# Bacterial defense and phage counter-defense lead to coexistence in a modeled ecosystem

**DOI:** 10.1101/2024.07.17.603905

**Authors:** Ofer Kimchi, Yigal Meir, Ned S. Wingreen

## Abstract

Bacteria have evolved many defenses against invading viruses (phage). Typically, each bacterium carries several defense systems, while each phage may carry multiple counter-defense systems. Despite the many bacterial defenses and phage counter-defenses, in most environments, bacteria and phage coexist, with neither driving the other to extinction. How is coexistence realized in the context of the bacteria/phage arms race, and how are the sizes of the bacterial immune and phage counter-immune repertoires determined in conditions of coexistence? Here we develop a simple mathematical model to consider the evolutionary and ecological dynamics of competing bacteria and phage with different immune/counter-immune repertoires. An analysis of our model reveals an ecologically stable fixed point exhibiting coexistence. This fixed point agrees with the experimental observation that each individual bacterium typically carries multiple defense systems, though fewer than the maximum number possible. However, in simulations, the populations typically remain dynamic, exhibiting chaotic fluctuations around this fixed point. We obtain quantitative predictions for the mean, amplitude, and timescale of these dynamics. Our results provide a framework for understanding the evolutionary and ecological dynamics of the bacteria/phage arms race, and demonstrate how bacteria/phage coexistence can stably arise from the coevolution of bacterial defense systems and phage counter-defense systems.

---

Bacteria and the viruses that infect them (phage) have been engaged in an arms race spanning eons. Each bacterium typically carries many defense systems to protect against phage [1]. Simultaneously, phage have evolved counter-defense systems that enable them to evade these bacterial defenses [2]. Both bacterial defense systems and phage counter-defense systems impose a fitness cost on the strains that carry them [3, 4]. In bacteria, for example, both metabolic costs associated with protein production [5] as well as inadvertent self-targeting [6] contribute to the fitness cost. In the absence of selection pressure from the opposing species, these systems are therefore quickly lost, over the timescale of a few generations [7, 8]. We sought to understand the co-evolution of these systems. What determines the size of a bacterial cell’s immune repertoire and a single phage’s counter-immunity repertoire?

One potential behavior of phage/bacteria community is that as a new phage strain invades, the bacteria may repel the invader by spreading defense systems through horizontal gene transfer [9]. This ‘pan-immunity hypothesis’ explains boom and bust cycles of phage and bacteria as arising from subsequent rounds of phage invasion and bacterial defense. In contrast, we wanted to study how persistent coexistence of phage and bacteria is realized within the context of the defense/counter-defense arms race.

We consider a mixture of bacterial strains with population densities *B*_*i*_ and phage strains with population densities *P*_*j*_. Each bacterial strain carries a subset of a total number *n*^tot^ possible defense systems, where we define *σ*_*id*_ = 1 if strain *i* carries defense system *d*, and 0 otherwise. The cost *γ*_*d*_ of each defense system reduces the growth rate *α*_*i*_ of a bacterial strain that carries it according to *α*_*i*_ = *g*(Σ_*d*_ *σ*_*id*_*γ*_*d*_) with a monotonically decreasing growth function *g*. Similarly, each phage strain carries a set of corresponding counter-defense systems, where *ρ*_*jd*_∈ 2 {0, 1}, such that phage *j* can infect bacteria *i* if it has a corresponding counter-defense system for each of bacteria *i*’s defense systems (Fig. 1) [10]. For phage, the cost of a counter-defense system, *β*_*d*_, is imposed on the phage burst size *b*_*j*_ +1 according to *b*_*j*_ + 1 = *h*(Σ _*d*_ *ρ*_*jd*_*β*_*d*_) for a monotonically-decreasing function *h*. We assume *g* and *h* are concave functions, but our results largely hold if they are convex (except in the *n*^tot^ → ∞ limit; see Supplementary Section S2). Bacterial and phage strain death rates are given by *μ*_*i*_ and *δ*_*j*_, respectively. Finally, we include steady immigration fluxes *λ*_*i*_ and *ν*_*j*_ for the bacteria and phage, respectively. The dynamics of this system are given by:

**Figure 1:**
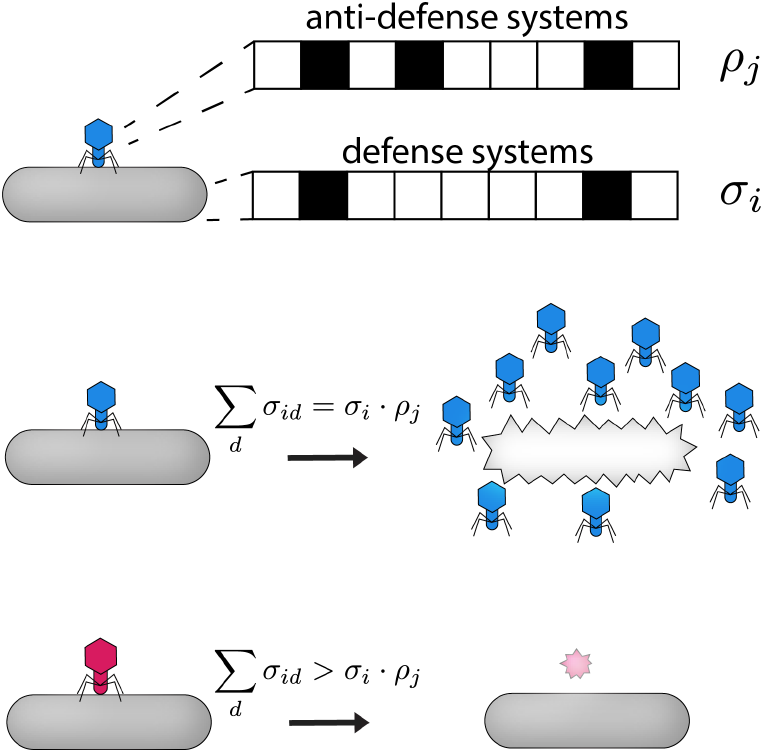
System overview. Pictorial representation of the model described by Eqs. (1). Each bacterial strain i has a set of defense systems *σ*_*i*_ and can only be infected by a phage strain j that has all the corresponding counter-defense systems *ρ*_*j*_ (see main text).

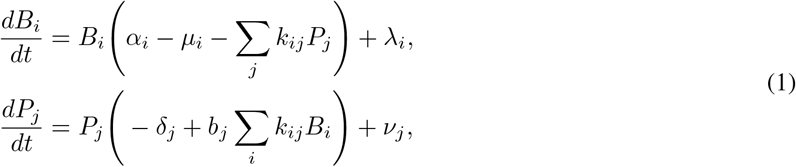

where the infection rate *k*_*ij*_ = *k* if phage *j* can infect bacteria *i* (i.e. if Σ_*d*_ *σ*_*id*_ = *σ*_*i*_ · *ρ*_*j*_) and 0 otherwise, and where we have neglected delays associated with phage reproduction [11, 12]. For simplicity, we focus on the symmetric case, where defense/counter-defense systems all have the same cost, i.e. *γ*_*d*_ = *γ* and *β*_*d*_ = *β*, and the immigration and death rates are strain-independent. In this case, it is useful to define a “largest positive-growth repertoire size” for bacteria and phage, labeled 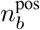 and 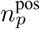, respectively, such that bacteria with 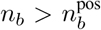 defense systems have a negative net growth rate *α* − *µ*, and phage with 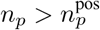 counter-defense systems have a negative net burst size, i.e. *b* < 0 (which takes into account the loss of the infecting phage particle).

To understand the expected behavior of this system, we first address Eqs. (1) analytically. For a system composed exclusively of bacteria with *n*_*b*_ defense systems and phage with *n*_*p*_ counter-defense systems, a dynamically stable fixed point can be found by solving 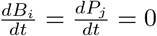. Due to symmetry, at this dynamical fixed point, all 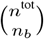 bacterial strains with *n*_*b*_ defense systems are present at equal population densities 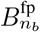, and all 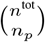 phage strains with *n*_*p*_ counter-defense systems are similarly present at equal population densities 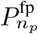. For any combination (*n*_*b*_, *n*_*p*_), such a dynamical fixed point exists. However, for certain combinations 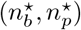, this dynamical fixed point will also be ecologically stable with respect to invasions by bacteria with 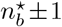 defense systems, and to phage with 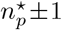 counter-defense systems, if the growth rates of such invading strains at the fixed point are negative. For large immigration rates *λ* and ***ν***, many such ecologically stable fixed points can exist, stabilized by the immigration flux. However, in realistic scenarios, immigration (as well as mutation and gene gain or loss) would be stochastic and would typically involve only small “incursions”; we therefore focus on ecologically stable fixed points in the limit *λ, ν* → 0. These ecologically stable fixed points 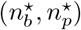 remain unchanged for small finite values of *λ* and *ν*, aside from minor shifts in their corresponding dynamical fixed-point population densities 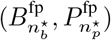. The conditions for such an ecologically stable fixed point are given by (see Supplementary Section S1)

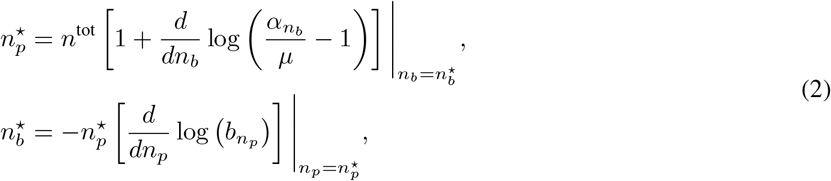

where the derivatives approximate a finite difference. Interestingly, Eqs. (2) predict that for small or zero immigration, the infection rate *k* and the phage death rate *δ* have no effect on the fixed-point number of bacterial defense or phage counter-defense systems per strain.

We find that Eqs. (2) typically predict that 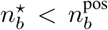, and similarly that 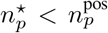 (Fig. 2a). In other words, at the ecologically stable fixed point, bacteria carry fewer than the maximal number of defense systems allowed by a positive growth rate, and similarly phage carry fewer than the maximal number of counter-defense systems.

**Figure 2:**
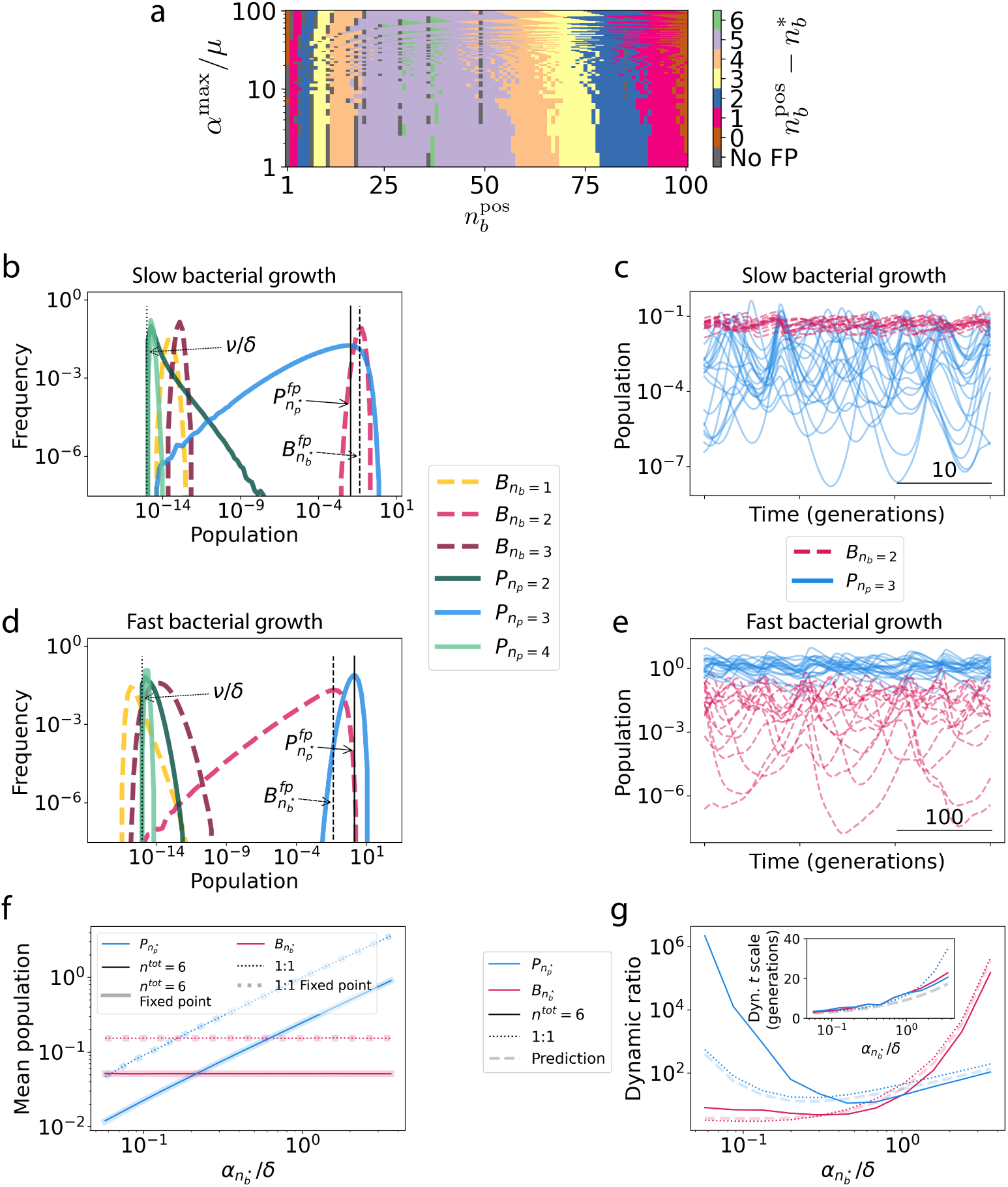
Fixed-point and dynamical simulation results. **a**, Predicted ecological fixed point for *n*^tot^ = 110 total defense systems, with 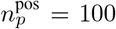 maximum counter-defense systems per phage strain. Distance between predicted ecological fixed point 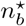 and maximal number of defense systems per bacterial strain, 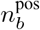, is shown as a function of 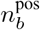 and the ratio between the growth rate of bacteria with no defense systems, *α*^max^, and the death rate of bacteria, *µ*. Gray points represent parameters for which no ecologically stable fixed point is predicted. **b**, Representative simulation results for an *n*^tot^ = 6 system with *α*^max^/*δ* = 0.1 and 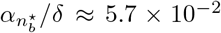, displaying the histogram of population densities of bacterial and phage strains with different numbers of defense/counter-defense systems per strain. **c**, Population dynamics of bacteria with 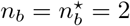 defense systems each (red) and phage with 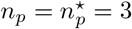 counter-defense systems each (blue) in the *n*^tot^ = 6 simulation of panel (b). **d, e** Representative simulation results as in (b), (c), but with bacterial growth rate increased to *α*^max^/*δ* = 10 (and 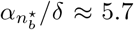). **f**, Dependence of mean population densities for *n*^tot^ = 6 system (solid curves), *n*^tot^ = 0 system (dotted curves), and analytical dynamical fixed-point prediction from Eqs. (1) (dashed curves). **g**, Dependence of dynamic ratio (main panel) and timescale of dynamics (inset) for *n*^tot^ = 6 system (solid curves), *n*^tot^ = 0 system (dotted curves), and analytical prediction (Eqs. (3); dashed curves). Here, initial conditions were kept constant independent of x-axis parameter variation; see Supplement for further discussion.

To explore the behavior of the system more fully, we turn to dynamical simulations with small equal immigration fluxes of all strains (see details in Supplementary Section S3). We consider a system with *n*^tot^ = 6 possible defense/counter-defense systems, with parameters such that 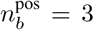 and 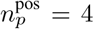, and for which Eqs. (2) predict the existence of an ecologically stable fixed point with 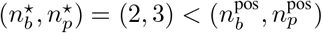. At the predicted ecologically stable fixed point, both bacterial growth rate and phage burst size are approximately half of their maximum values: 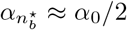 and 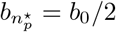. In agreement with our analytical results, we find that the system evolves to a state where all bacterial strains that are present at high population densities have 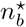 defense systems each, and all phage at high population densities have 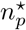 counter-defense systems each (Fig. 2b). In the absence of immigration (*λ* = *ν* = 0) only these strains survive. Moreover, the analytically predicted dynamical fixed-point population densities predict well the time-averaged population densities of these strains (Fig. 2b, vertical lines). However, we do not find that the system reaches the corresponding dynamical fixed point. Rather, the system displays persistent chaotic dynamics, even at long times (Fig. 2c; Lyapunov exponent = 0.08, see Supplementary Section S3) [13].

Notably, the chaotic population fluctuations enable coexistence to be maintained even when the number of phage “predator” strains,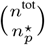, exceeds the number of bacterial “prey” strains 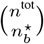 [12, 14]. For different parameter choices, the chaotic fluctuations enabling this coexistence may either be largest in the phage population (as in Fig. 2b,c) or largest in the bacterial population (as in Fig. 2d,e). This variation in the dynamics comes with no apparent effect on the coexistence itself: we find that all bacteria with 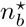 defense systems and all phage with 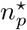 counter-defense systems coexist in all cases we examined, even in the absence of immigration, *λ* = *ν* = 0, with the system initialized to include only bacterial strains with 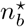 defense systems and phage with 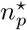 counter-defense systems. Sufficient parameter heterogeneity can lead some strains to go extinct, but we found no extinction events within 10^6^ generations for moderate amounts of parameter heterogeneity (𝒪 (10^−5^)).

We sought to understand how the system dynamics depend on parameters. While the average population densities are well predicted by the dynamical fixed-point population densities (Fig. 2f), the dynamics cannot be predicted by the fixed-point analysis alone. Helpfully, we find that a quantitatively similar parameter dependence of dynamics occurs in the *n*^tot^ = 0 case which corresponds to a minimal system of one bacterial strain and one phage strain, and which results in oscillatory dynamics (hereafter referred to as the 1:1 case). The dynamic ratios of the population densities in the 1:1 case scale with parameters according to (see derivation in Supplementary Section S4):

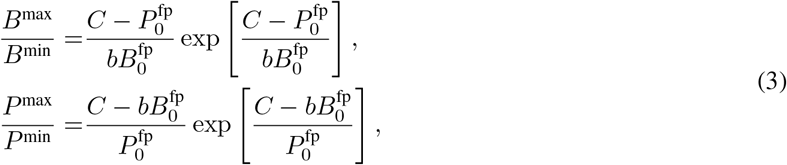

where *C* is a time-independent conserved quantity:

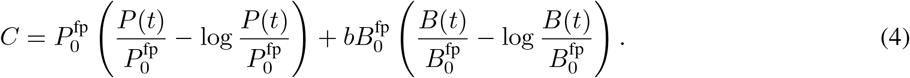

The parameter-dependent dynamic ratio, as well as the timescale of the dynamics (predicted by linear stability analysis; see Supplementary Section S4), are both in quantitative agreement between the *n*^tot^ = 6 and 1:1 cases (Fig. 2g). The growth of the dynamic ratio of bacterial population densities for large *α* can be understood from Eqs. (3) to result from the exponential term 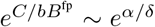 (see Supplementary Section S4). The same scaling behavior applies to the growth of the dynamic ratio of phage population densities for small *α*: for *α* ≪ *δ*, the exponential term e^*δ*/(*α*−*µ*)^ dominates the dynamic ratio of *P*. The crossover between the regime of dominant bacterial dynamics and that of dominant phage dynamics occurs when *P*^fp^ = *bB*^fp^, i.e. when *α* − *µ* = *δ* (see Supplementary Section S4). Using the analytically tractable 1:1 system as a guide, we are thus able to predict the dynamic ratios and timescales of the parameter-dependent chaotic dynamics for more complex cases with multiple bacterial and phage strains.

We note that our model makes several simplifying assumptions. While we have considered all defense systems to be qualitatively interchangeable, defense systems in nature operate through different mechanisms and may have qualitatively different effects on both the growth cost to the bacterium and on the success or failure of the invading phage (and similarly for phage counter-defense systems) [3]. Interactions among defense systems may also change their efficacies [15, 16]. We have also focused exclusively on obligate lytic (virulent) phage, neglecting temperate phage as well as other alternative phage infection strategies [17]. While temperate phage may qualitatively affect the behavior of many phage-bacteria interactions [18], here, the effect of super-immunity exclusion (i.e. that lysogens are immune to further infection by phage of the same strain that lysogenized them) may be considered as a special case of a defense system. Other phage infection strategies may also have only minor effects on our results; for example, chronic infections wherein phage reproduce and exit the cell without cell lysis may be considered as a modification of the burst size b. Furthermore, although we have here assumed well-mixed populations, spatial organization likely affects both the dynamics of phage/bacteria competition and their coexistence, particularly in non-aquatic environments [19]. In this regard, abortive infection defenses will be a fruitful topic for future work. Finally, while we have focused here on steady immigration fluxes, a variant of the model described by Eqs. (1), without immigration fluxes but allowing the bacteria and phage to stochastically gain, lose, and exchange systems with one another through mutations and horizontal gene transfer, yields qualitatively similar results (see Supplementary Section S3).

In summary, we have developed a model for the dynamics of competing phage and bacteria with different sets of defense and counter-defense systems. A fixed-point analysis of the model (confirmed by dynamical simulations) indicates that phage and bacteria typically evolve to have more than one and less than the maximum number of defense/counter-defense systems in each strain. This qualitative behavior has been observed in nature, and has previously been explained by the pan-immunity hypothesis, which argues that invading phage can be driven to extinction as long as some bacteria within a community are immune to the invading phage, and that this immunity can be conferred to other bacteria through horizontal gene transfer [9]. In contrast, we find that within our model, large (but non-maximal) immunity and counter-immunity repertoires emerge naturally and enable the coexistence of phage and bacteria. This coexistence is manifested in persistent chaotic dynamics, with the mean, amplitude, and timescale of these dynamics well predicted by an analysis based on the competition between a single bacterial and a single phage strain. Although the ecological fixed point of the system 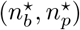 depends on the details of the growth costs of defense and counter-defense systems through 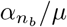 and 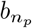 (Eq. (2)), we find that the qualitative dynamics depend only on the ratio *α*/*δ*, so long as immigration is minimal and *µ* is small enough that bacteria mostly die due to phage predation. Thus, there are two main regimes of system behavior: slow bacterial growth rate compared to phage death rate (Fig. 2b,c); and fast bacterial growth rate compared to phage death rate (Fig. 2d,e). Finally, although we have focused our analytical analysis on symmetric systems, our simulations demonstrate that even non-symmetric systems with modest amounts of heterogeneity are not limited by the principle of competitive exclusion in our model. Thus, we find that the chaotic dynamics of the system enable the coexistence of more phage (predator) strains than bacterial (prey) strains, exceeding the biodiversity predicted by other frameworks such as “kill-the-winner” [20, 21].

The discovery that bacteria typically contain multiple coexisting defense systems (and phage contain multiple counter-defense systems) has raised many questions. Chief among these, what controls the number and type of defense and counter-defense systems in a particular bacterium or phage? One possibility is that these systems are controlled by happenstance, with horizontal gene transfer mediating random gains and losses of systems. Alternatively, our simple model suggests that considering states of evolutionary stability–albeit with fluctuations about these states–may provide a helpful guiding perspective.

## Acknowledgements

We thank Wenping Cui, Simon Levin, and Pankaj Mehta for useful discussions. This work was supported in part by grant number DAF2024-342781 from the Chan Zuckerberg Initiative DAF, an advised fund of Silicon Valley Community Foundation, by the National Science Foundation through the Center for the Physics of Biological Function (PHY-1734030), and by the Peter B. Lewis ‘55 Lewis-Sigler Institute/Genomics Fund through the Lewis-Sigler Institute of Integrative Genomics at Princeton University (O.K.). This work was performed in part at Aspen Center for Physics, which is supported by National Science Foundation grant PHY-1607611.

## Supplement

### S1 Deriving Eqs. (2)

To derive Eqs. (2), we start from Eqs. (1), making the simplifications discussed in the main text, and assuming that the immigration terms *λ* and *ν* are negligible for the purpose of the following analyses. We consider a system composed exclusively of bacteria with *n*_*b*_ defense systems and phage with *n*_*p*_ counter-defense systems. At a dynamical fixed point of such a system, due to symmetry, all 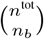 bacterial strains with *n*_*b*_ defense systems are present at equal population densities 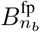, and all 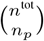 phage strains with *n*_*p*_ counter-defense systems are present at equal population densities 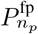. This dynamical fixed point is given by

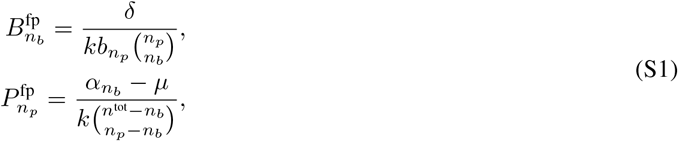

where 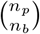 is the number of bacterial strains each phage strain can infect, 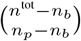 and is the number of phage strains that can infect each bacterial strain.

This dynamical fixed point will further be an ecological fixed point if it is stable to invasions by bacteria with a different number of defense systems, or to phage with a different number of counter-defense systems. This will occur if the growth rates of such invading strains at the dynamical fixed point are negative. We refer to such an ecologically stable fixed point as 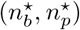. The growth rate of an invading bacterial strain with 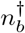 defense systems at the ecological fixed point would be

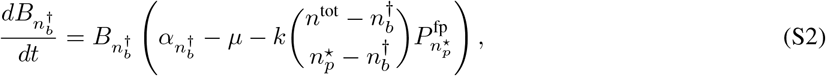

where 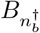 is the population of the invading bacterial strain. Similarly, the growth rate of an invading phage strain with 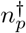 counter-defense systems would be

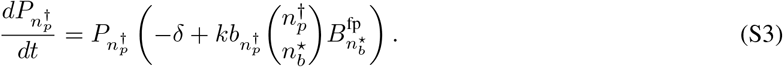

These terms are negative for invading bacterial strains with 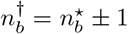 defense systems, and for invading phage strains with 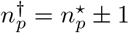 counter-defense systems, when

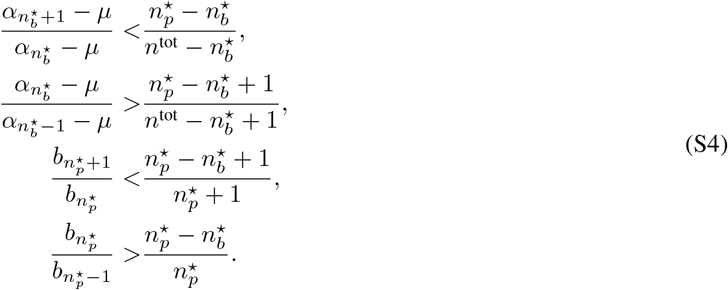

By taking the natural logarithm and substituting derivatives for differences, we arrive at Eqs. (2).

### S2 Ecological fixed point behavior as *n*^tot^ → ∞

The behavior of the ecological fixed point defined by Eqs. (2) as *n*^tot^ → ∞ depends on how 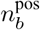 and 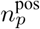 scale with n. Since ecological stability requires 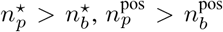 must hold to avoid the trivial outcome with no phage present. This inequality is biologically reasonable given the complexity required of defense systems compared to the relative simplicity of counter-defense systems; as an example, consider the immensely complex Type 1-F CRISPR-Cas system which can be evaded by phage that express a single short RNA molecule [S1]. As *n*^tot^ → ∞, there are therefore three possibilities: 1) 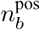 and 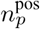are both intensive; 2) 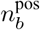 and 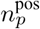 are both extensive; 3) 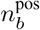is intensive while 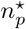 is extensive. We find that in cases (1) and (2), 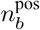 and 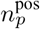 are approximately equal to 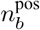 and 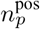, respectively, as n ^tot^ → ∞. However, surprisingly, in case (3) and for concave cost functions *g* and *h*, 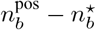 and 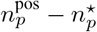both grow with *n*^tot^. In this case, each individual bacterium optimally carries a subset of possible defense systems, even though it could carry far more and still continue to grow (and similarly for phage).

### S3 Details of dynamical simulations

We use the following parameters in our simulations, with time units such that *δ* = 1 and population density units such that *k* = 1: *µ* = 10^−2^; *λ* = *ν* = 10^−15^; *b*^max^ = 15. Bacterial growth rate *α* is varied as described in the main text; all other parameters are kept constant throughout. The bacterial death rate *µ* = 10^−2^ was chosen to be much smaller than the phage death rate *δ* so that we can explore the *α* < *δ* regime while maintaining *µ* ≪*α*, such that bacterial populations are primarily limited by phage predation. For example, for 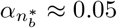 as in Fig. 2b,c, the rate of bacterial death due to phage predation at the dynamical fixed point, 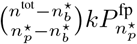, is approximately 5× larger than *µ*; for 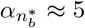 as in Fig. 2d,e, it is approximately 500× larger. The particular values of *λ* and *ν* matter very little as long as they are in the regime of slow immigration. (As described in the main text, the opposite regime where immigration is substantial is qualitatively different because immigration stabilizes the populations of phage and bacteria strains which would otherwise go extinct). Finally, the particular value of the phage burst size b has very little effect on our results since the ecological fixed point depends on 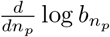(and is therefore unchanged when b is modified by a multiplicative factor; Eq. (2)) and the qualitative dynamical behavior is mostly determined by *α*/*δ* as described in Section S4. Natural phage burst sizes are typically of 𝒪 (100) phage particles per burst, but also are accompanied by a sizeable time delay between phage infection and lysis. Given that our model neglects this time delay, the effective burst size must be decreased to reproduce overall phage proliferation rates. *b*^max^ = 15 was therefore chosen to correspond to a burst size of ∼ 200 for a system with a typical lysis time [S2].

We implement concave cost functions 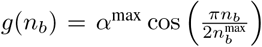 and 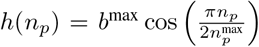, where we have set *γ*_*d*_=1 for all defense systems and *β*_*d*_=1 for all counter-defense systems.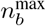 sets the maximum number of defense systems bacteria can have before their growth rate *α* reaches zero, and is somewhat larger than 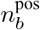 which is determined by the net growth rate *α* − *µ* reaching zero (and similarly for 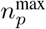 for phage). Strains with more systems than the maximum don’t grow: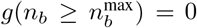, and similarly,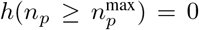. In our simulations, we set 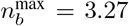, and 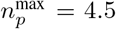, as these values yielded 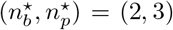 and 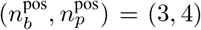 for our parameter choices.

In all simulations, the growth rates of individual bacterial strains *α*_*i*_, the burst sizes of individual phage strains b_*j*_, and the infection rates *k*_*ij*_ were all varied at order 10^−5^. Specifically, these terms were multiplied by 1 + 10^−5^ × (2*r* − 1) where r is a random number uniformly distributed between 0 and 1, chosen randomly and independently for each strain or strain-strain interaction. Thus, small amounts of parameter heterogeneity are added to the system; for example, values of k ultimately range from (1 − 10^−5^) to (1 + 10^−5^).

We ran each simulation for 10^7^ time units (equivalent to 1.4 × 10^6^ bacterial generations for *α*/*µ* = 10, or to more generations for larger *α*).

We also ran simulations initializing the system with only bacterial strains for which 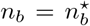 and phage strains with 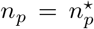, and setting the immigration fluxes to zero, *λ* = *ν* = 0. We found qualitatively and quantitatively similar results to those displayed in Fig. 2b-e, and observed no phage or bacterial strains going extinct over the course of these simulations. The simulations in Fig. 2f-g were performed in this manner.

To measure the Lyapunov exponent, we initialized two trajectories using the parameters of Fig. 2b,c, with initial values of phage populations ∼ 10^−14^ higher in one trajectory than the other. Specifically, the initial values in the second simulation were equal to those of the first, plus 10^−14^ × (2*r* − 1), where *r* is a uniformly distributed random number between 0 and 1 chosen independently for each phage strain. We then measured the distance between the two trajectories as 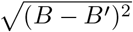 where B is the total bacterial population. This value grows exponentially until ∼ 400 generations, at which point it begins fluctuating between 10^−5^ and 10^0^. This behavior of the distance between trajectories initialized nearby (namely, exponential growth, followed by fluctuations around a fixed value) is typical of chaotic systems. The Lyapunov exponent is defined as the slope of the exponential growth segment. To measure this, the logarithm of the distance was fit to a line (or equivalently, the distance was fit to an exponential). This analysis was repeated ten times for different instantiations of *r*, with measurements of the Lyapunov exponent having a mean of 0.0822 and a standard deviation of 0.0014.

In Fig. 2g, we needed to measure the dynamic ratio in a consistent way for both chaotic and oscillatory systems. To do this, first, we identified the local peaks and troughs (i.e. where the derivatives of the population densities change sign). For oscillatory systems, the peaks are all at (very nearly) equal values, as are the troughs. For chaotic systems, this is not the case. In order to measure the peaks and troughs in a manner that does not change much between different instantiations of the chaotic dynamics, we therefore defined the dynamic ratio as the ratio of the 90^th^ percentile of the local population density peaks to the 10^th^ percentile of the local population density troughs. The dynamic timescale was measured by taking the median of the time between local maxima and the median of the time between local minima, and averaging these two medians. For oscillatory systems, this measures the period of oscillations.

The simulations in Fig. 2f-g were initiated at *B*_*i*_ = 0.06 × (1 + *r*) and *P*_*j*_ = 0.06 × (1 + *r*) where r is a random number uniformly distributed between 0 and 1, chosen independently for each bacterial and each phage strain. Qualitatively similar results were obtained for simulations initiated at 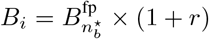 and 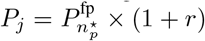 i.e. where the initial conditions depend on parameter values through the dynamical fixed-point population densities. Finally, we developed a variant of these simulations to take into account the stochasticity of evolutionary dynamics.

Rather than initiating the system with all strains present, we initialized the system with only one bacterial strain (with one defense system), and one phage strain (with the corresponding counter-defense system). After every 5 timepoints of simulation, we performed a mutation step. In this mutation step, new bacterial or phage strains could be created, by either gaining or losing defense or counter-defense systems. First, we choose whether to mutate the bacteria (with probability *p*) or the phage (with probability 1 − *p*); we chose *p* = 10^−2^ to approximate the challenge bacteria face in constructing new defense systems as opposed to the relative simplicity of phage counter-defense systems, as discussed in the previous section. Next, we select the strain to mutate, proportionally to its population density at the time of the mutation step. We then determine whether the mutation will be the (a) gain or (b) loss of defense (or counter-defense) systems, each occurring with probability 1/2. Finally, all possible single mutants either adding or removing a defense or counter-defense system (depending on which was selected) from the selected strain are added to the system at a small initial population (10^−15^), or if they were already present, their population is increased by the same small amount. These simulations resulted in the same qualitative behavior as the constant immigration rate simulations discussed in the main text, evolving towards the ecologically stable fixed point at the system level, and exhibiting chaotic dynamics at the population level.

### S4 Dynamical analysis

To understand the dynamical behavior of the system, we turn to the *n*^tot^ = 0 case, which we refer to as the 1:1 case since it involves a single bacterial strain and a single phage strain. In the main text, we show that certain aspects of the dynamical behavior of the 1:1 case closely parallel those of systems with *n*^tot^ > 0. In this section, we quantitatively analyze the 1:1 case.

First, we perform linear stability analysis to describe the behavior of the system near the dynamical fixed point. The essential element of linear stability analysis is the calculation of the eigenvalues of the Jacobian matrix at the dynamical fixed point (Eqs. (S1)). For the 1:1 case, the eigenvalues are 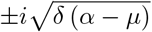. That these are imaginary indicates that the 1:1 case undergoes continued oscillations. The period of these oscillations is given by 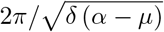. This prediction is plotted as a dashed curve in the inset to Fig. 2g.

Predicting the amplitude of oscillations is less straightforward, and to our knowledge no generic method enables this prediction. Linear stability analysis provides no information regarding the amplitude of oscillations. To address this challenge, we recognize that there is a quantity C satisfying dC/dt = 0. In general, this quantity is given by

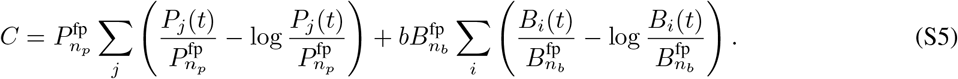

For the 1:1 case, this quantity simplifies to

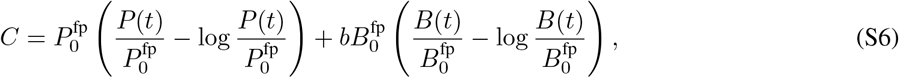

where 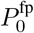 and 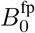 are given by Eqs. (S1) with *n*_*p*_ = *n*_*b*_ = *n*^tot^ = 0.

Because *C* is a constant in time, it will also be a constant when *P* or *B* are at an extremum. Solving for *B* at either extremum of *P* (i.e. where *dP*/*dt* = 0) yields 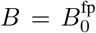. Similarly, solving for *P* at either extremum of *B* yields 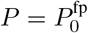. Thus, we find that

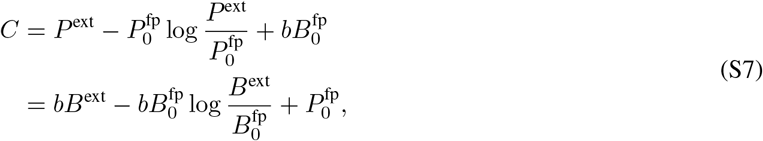

where *B*^ext^ represents the value of *B*(*t*) at its maximum or minimum, and similarly for P^ext^.

We then estimate the values of *P*^min^, *P*^max^, *B*^min^, and *B*^max^. For the minimum values, we treat the linear terms (e.g. *P*^ext^) as negligible compared to the logarithmic terms; for the maximum values, we treat the logarthmic terms as negligible. These approximations yield

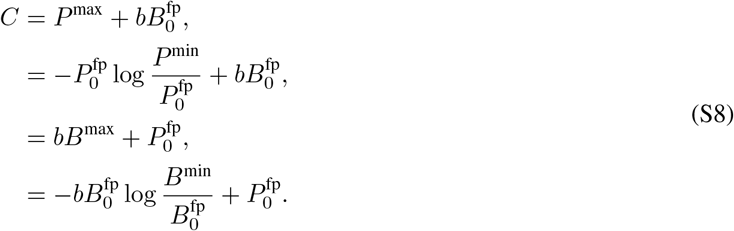

Solving for the extrema and simplifying, we find that the dynamic ratios of *B* and of *P* are given by Eqs. (3), reprinted here:

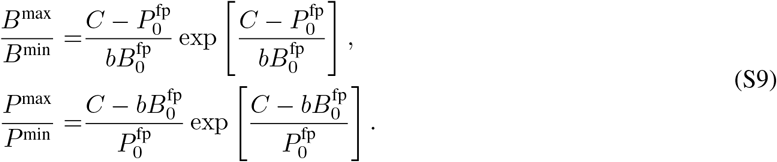

To understand the growth of the dynamic ratio of *B* for large *α* and of *P* for small *α*, we first recognize that the positivity of the dynamic ratios implies 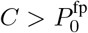 and 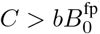, so that the dynamic ratios can be approximated as

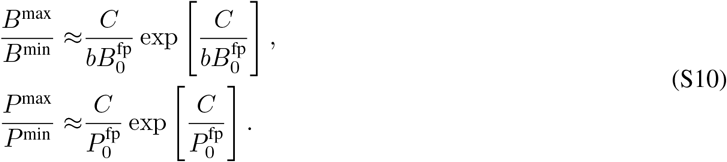

Substituting in from Eqs. (S8), we find

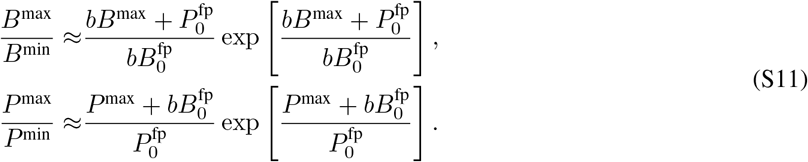

Finally, substituting in for the dynamical fixed point values (Eqs. (S1)), we arrive at

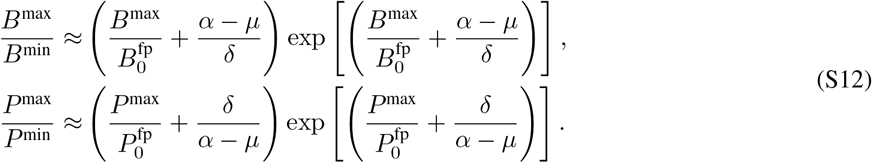

For large *α* (i.e. *α*≫ *δ*), the dynamic ratio of B is therefore dominated by e^*α*/*δ*^. Similarly, for small *α* (i.e. *α*−*µ* ≪ *δ*), the dynamic ratio of *P* is dominated by e^*δ*/(*α*−*µ*)^.

To find the crossover between the two dynamic regimes (one where the dynamic ratio of B is large, and the other where the dynamic ratio of P is large), we solve for

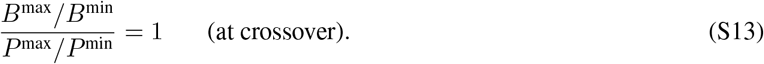

Starting from Eq. (S10), this can be approximated as

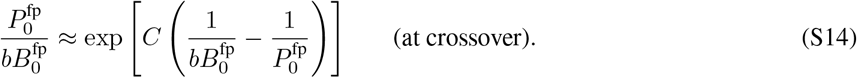

While the precise value of C depends on the particular initial conditions chosen, it can be estimated by its value at the fixed point, 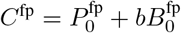. This yields

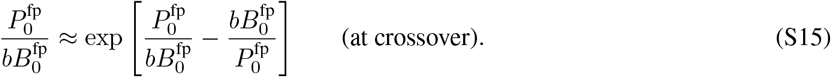

The equation x = exp [*x* − *x*^−1^]is solved by *x* = 1. Therefore, the crossover occurs at

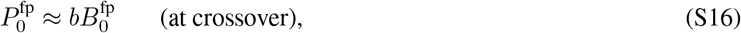

or, substituting in for the dynamical fixed point values (Eqs. (S1)),

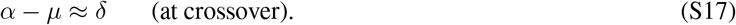

## References

[1] Héloïse Georjon and Aude Bernheim. The highly diverse antiphage defence systems of bacteria. Nature Reviews Microbiology, 21(10):686–700, 2023.

[2] Khalimat Murtazalieva, Andre Mu, Aleksandra Petrovskaya, and Robert D. Finn. The growing repertoire of phage anti-defence systems. Trends in Microbiology, xx(xx):1–17, 2024.

[3] Hannah G. Hampton, Bridget N.J. Watson, and Peter C. Fineran. The arms race between bacteria and their phage foes. Nature, 577(7790):327–336, 2020.

[4] Sriram Srikant, Chantal K. Guegler, and Michael T. Laub. The evolution of a counter-defense mechanism in a virus constrains its host range. eLife, 11:1–25, 2022.

[5] Pedro F. Vale, Guillaume Lafforgue, Francois Gatchitch, Rozenn Gardan, Sylvain Moineau, and Sylvain Gandon. Costs of CRISPR-Cas-mediated resistance in Streptococcus thermophilus. Proceedings of the Royal Society B: Biological Sciences, 282(1812), 2015.

[6] Maroš Pleška, Long Qian, Reiko Okura, Tobias Bergmiller, Yuichi Wakamoto, Edo Kussell, and Calin C. Guet. Bacterial autoimmunity due to a restriction-modification system. Current Biology, 26(3):404–409, 2016.

[7] Pedro Gómez and Angus Buckling. Bacteria-phage antagonistic coevolution in soil. Science, 332(6025):106–109, 2011.

[8] Joshua M. Borin, Justin J. Lee, Adriana Lucia-Sanz, Krista R. Gerbino, Joshua S. Weitz, and Justin R. Meyer. Rapid bacteria-phage coevolution drives the emergence of multiscale networks. Science, 382(6671):674–678, 2023.

[9] Aude Bernheim and Rotem Sorek. The pan-immune system of bacteria: antiviral defence as a community resource. Nature Reviews Microbiology, 18(2):113–119, 2020.

[10] Simon A. Levin, Lee A. Segel, and Frederick R. Adler. Diffuse coevolution in plant-herbivore communities. Theoretical Population Biology, 37(1):171–191, 1990.

[11] Rasmus Skytte Eriksen, Namiko Mitarai, and Kim Sneppen. Sustainability of spatially distributed bacteria-phage systems. Scientific reports, 10(1):3154, 2020.

[12] Ofer Kimchi, Yigal Meir, and Ned S Wingreen. Lytic and temperate phage naturally coexist in a dynamic population model. The ISME Journal, 18(May):1–5, 2024.

[13] Edward H. Kerner. A statistical mechanics of interacting biological species. The Bulletin of Mathematical Biophysics, 19(2):121–146, 1957.

[14] Jef Huisman and Franz J. Weissing. Biodiversity of plankton by species oscilations and chaos. Nature, 402:407–410, 1999.

[15] Clemente F. Arias, Francisco J. Acosta, Federica Bertocchini, Miguel A. Herrero, and Cristina Fernández-Arias. The coordination of anti-phage immunity mechanisms in bacterial cells. Nature Communications, 13(1):1–11, 2022.

[16] Yi Wu, Sofya K. Garushyants, Anne van den Hurk, Cristian Aparicio-Maldonado, Simran Krishnakant Kushwaha, Claire M. King, Yaqing Ou, Thomas C. Todeschini, Martha R.J. Clokie, Andrew D. Millard, Yilmaz Emre Gençay, Eugene V. Koonin, and Franklin L. Nobrega. Bacterial defense systems exhibit synergistic anti-phage activity. Cell Host and Microbe, 32(4):557–572.e6, 2024.

[17] Sari Mäntynen, Elina Laanto, Hanna M. Oksanen, Minna M. Poranen, and Samuel L. Díaz-Muñoz. Black box of phage-bacterium interactions: Exploring alternative phage infection strategies. Open Biology, 11(9):8–10, 2021.

[18] Michael G. Cortes, Yiruo Lin, Lanying Zeng, and Gábor Balázsi. From Bench to Keyboard and Back Again: A Brief History of Lambda Phage Modeling. Annual Review of Biophysics, 50:117–134, 2021.

[19] Samuele Testa, Sarah Berger, Philippe Piccardi, Frank Oechslin, Grégory Resch, and Sara Mitri. Spatial structure affects phage efficacy in infecting dual-strain biofilms of Pseudomonas aeruginosa. Communications Biology, 2(1):1–12, 2019.

[20] T. F. Thingstad. Elements of a theory for the mechanisms controlling abundance, diversity, and biogeochemical role of lytic bacterial viruses in aquatic systems. Limnology and Oceanography, 45(6):1320–1328, 2000.

[21] Anastasios Marantos, Namiko Mitarai, and Kim Sneppen. From kill the winner to eliminate the winner in open phage-bacteria systems. PLoS Computational Biology, 18(8):1–15, 2022.

## Supplemental References

[S1] Sarah Camara-Wilpert, David Mayo-Muñoz, Jakob Russel, Robert D. Fagerlund, Jonas S. Madsen, Peter C. Fineran, Søren J. Sørensen, and Rafael Pinilla-Redondo. Bacteriophages suppress CRISPR–Cas immunity using RNA-based anti-CRISPRs. Nature, 623(7987):601–607, 2023.

[S2] Ofer Kimchi, Yigal Meir, and Ned S Wingreen. Lytic and temperate phage naturally coexist in a dynamic population model. The ISME Journal, 18(May):1–5, 2024.

